# Biomechanical simplification of the motor control of whisking

**DOI:** 10.1101/2025.06.21.660818

**Authors:** Chris S. Bresee, Yifu Luo, Jasmine L. Alade’Fa, Megan E. Black, Kevin J. Kleczka, Nicholas E. Bush, Kevin Zhang, Mitra J. Hartmann

## Abstract

Animal nervous systems must coordinate the sequence and timing of numerous muscles − a challenging control problem. The challenge is particularly acute for highly mobile sensing structures with many degrees of freedom, such as eyes, pinnae, hands, forepaws, and whiskers, because these low-mass, distal sensors require complex muscle coordination. This work examines how the geometry of the rat whisker array simplifies coordination required for “whisking” behavior ^1-3^. During whisking, 33 intrinsic (“sling”) muscles are the primary drivers ^4-12^ of the rapid, rhythmic protractions of the large mystacial vibrissae (whiskers), which vary more than sixfold in length and threefold in base diameter ^13-16^. Although whisking is a rhythmic, centrally-patterned behavior ^17-24^, rodents can change the position, shape, and size of the whisker array, indicating considerable voluntary control ^25-34^. To begin quantifying how the array’s biomechanics contribute to whisking movements, we used three-dimensional anatomical reconstructions of follicle and sling muscle geometry to simulate the movement resulting from a “uniform motor command,” defined as equal firing rates across all sling muscle motor neurons. This simulation provides a baseline profile of protraction under anatomically realistic but uniformly driven conditions. It does not isolate neural from biomechanical contributions but helps identify deviations that suggest active control. Simulations reveal that all follicles rotate through approximately equal angles, regardless of size. The angular fanning of the whiskers at their bases increases monotonically throughout protraction, while maximum distance between whisker tips occurs at approximately 90% of resting muscle length, after which whisker tips converge and sensing resolution increases monotonically.

## 2. Results and Discussion

### 2.1. The three dimensional (3D) geometry of follicle motion is defined by a triangle formed by follicle lever length, sling muscle length, and the distance between follicles

Whisking behavior is critical for rodents, enabling them to tactually explore their surroundings and discern object size, shape, texture, and orientation ^35,36^. However, whisking movements exhibit considerable variability, even on a whisk-to-whisk basis ^26,27,29,32^. Understanding the origins of this variability, and the extent to which it arises from neural control or from the biomechanics of the whisker array, requires careful anatomical and functional analysis.

As illustrated in Fig. 1A, the follicles and whiskers on the rat’s mystacial pad are arranged in rows (A through E) and columns (1 through 6). The most caudal whiskers are denoted alpha through delta. During whisking, intrinsic (“sling”) muscles govern whisker protractions ^4-12^, and the minimum actuation unit is a row-wise pair of adjacent follicles ^8^. Within each row, each sling muscle originates at the apex of a caudal follicle and wraps around the base of its adjacent rostral follicle (Fig. 1B) ^4,8-10,12,37,38^. During sling muscle contraction, the skin acts as a fulcrum, causing the caudal follicle to rotate and protract the whisker. Four parameters thus define basic protraction geometry: gap *G*, lever length *L*, muscle arm length *m*, and follicle circumference *C*. The gap is the straight-line distance between a follicle apex and that of its caudal neighbor; the lever length is the distance from the apex to the sling muscle attachment; the muscle arm length is the distance from the sling muscle attachment point to the apex of its caudal neighbor. The circumference (*C*) is measured at the muscle attachment point. The triangle formed by *G, L*, and *m* is termed the “*GLm* triangle.”

**Figure 1.**
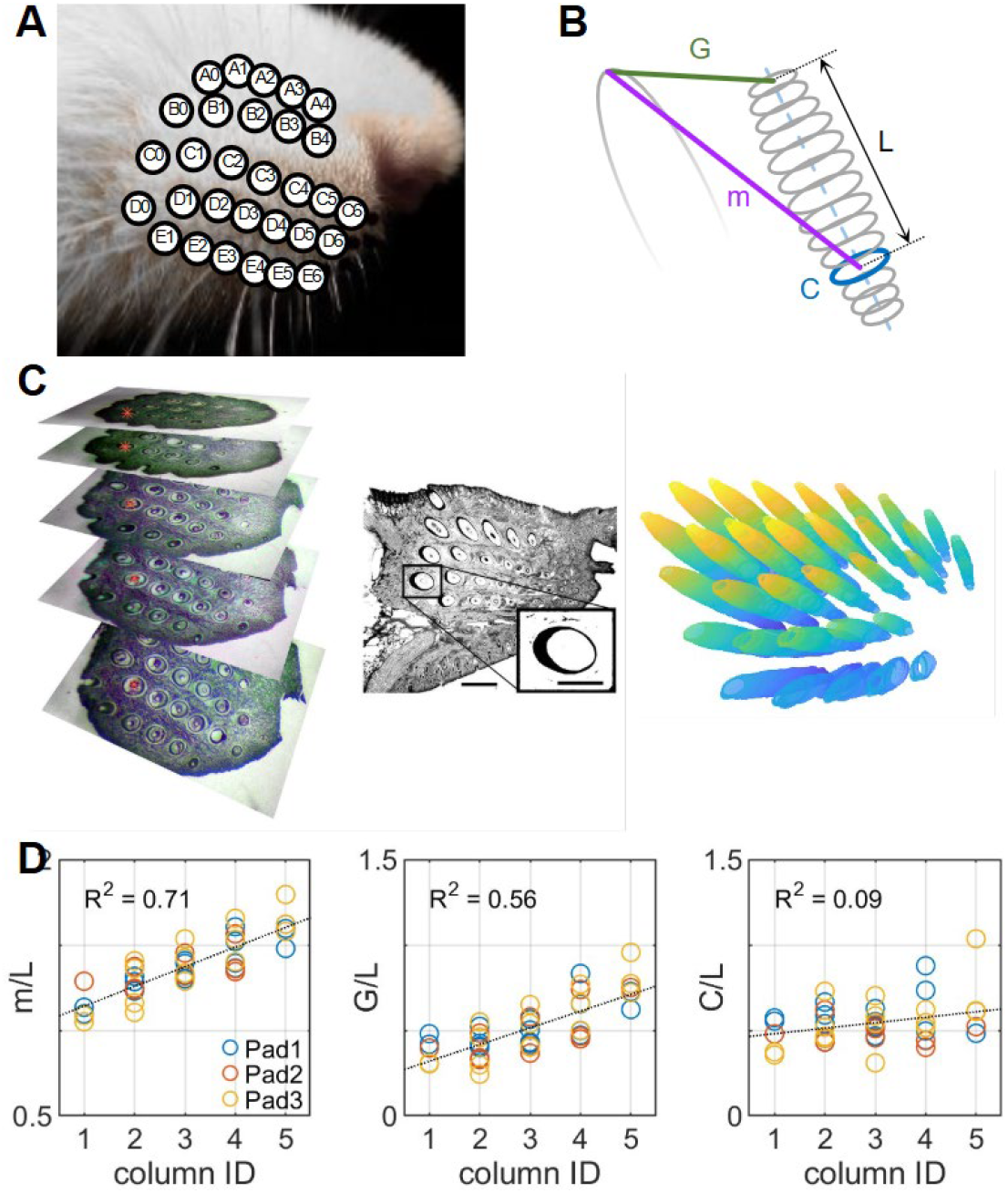
Quantification of follicle and sling muscle geometry. **(A)** Arrangement of mystacial vibrissae. **(B)** The parameters *G, L, m*, and *C* determine follicle rotation; the first three parameters form the “*GLm* triangle.” **(C)** Mystacial pads were dissected, fixed, and sliced; the photomicrograph stack shows FG/PTOH stained sections. Expanded view shows a single follicle from a typical 8-bit grayscale image used for analysis. Red asterisks were manually added to the left panel to aid reader visualization of a single follicle. Right panel shows one view of the follicles reconstructed from Pad 1. **(D)** The ratios *m*/*L* and *G*/*L* vary with column position, while *C*/*L* does not. The three colors represent data from three mystacial pads. Fig. 1A adapted from Belli et al., 2018, published under a CCBY 4.0 license. Related to Table 1, Fig. S2, and Video S1.

To begin quantifying how the array’s biomechanics shape whisking movements, we simulated the movement consequences of a “uniform motor command,” defined as the motion that would result if all motor neurons controlling the sling muscles fired at the same rate. This simulation does not by itself disentangle relative contributions of neural versus biomechanical factors in whisking variability. Rather, it provides a baseline profile for whisker protraction under anatomically realistic, uniformly driven conditions; any observed deviations from this profile point to additional neural control.

To build the simulation, we reconstructed follicle and sling-muscle anatomy across the central mystacial pad, obtaining estimates for *G, L, m* and *C* for each whisker (*Methods*, Table S1). Some previous studies ^37^ have used a customized flattening technique to preserve relative whisker positions, while others have focused on anatomical analysis of a single follicle row ^39,40^. Here, we embedded and then sliced three mystacial pads from three adult, Long Evans rats, with the goal of maintaining the pads’ natural curvatures and quantifying 3D follicle positions and orientations in the pads’ central regions. In a set of control experiments, we performed measurements of sarcomere stretch to confirm that sling muscles (and thus follicles) were near rest (*Methods*; Fig. S1).

General procedures are illustrated from left to right in Fig. 1C; details are in *Methods*. Because sectioning was perpendicular to the pad, follicles were sliced at varying angles, requiring anatomical parameters to be computed rather than measured. Mystacial pads were dissected, fixed, sliced (either 20- or 50-micron sections), and then stained with Fast Green (FG; for collagen) and Mallory’s Phosphotungstic Acid Hematoxilin (PTAH; for muscles). After flatbed scanning, images were converted to grayscale, and follicle centers and cross-sections detected with standard image processing. Each follicle cross-section was fitted with an ellipse, and these ellipses were aligned, smoothed, and resliced to reconstruct all 3D follicle shapes. Muscle lengths were found by identifying, for each follicle pair within a row, the first section in which the sling muscle fully wrapped around the rostral follicle; the corresponding location in the resliced follicle was taken as the muscle attachment point.

A total of 96 follicles were reconstructed from the three pads. Follicles reconstructed from Pad 1 are visualized in the right panel of Fig. 1C and rotated in 3D in Video S1. After excluding poorly reconstructed follicles (see *Methods*), 43 follicles from 17 unique row-column identities were measured along with their sling muscle attachments. All follicles were located in the central pad region and are identified in Table S1. Consistent with previous studies ^39,40^, reconstructions revealed that the lower third of many follicles appeared to curve, likely due to reduced tissue stiffness at depth ^41^. Follicles were therefore linearly fit to match previously established 3D whisker emergence angles ^42^.

Analysis of follicle and muscle geometry, shown in Fig. 1D, revealed that the ratios *m*/*L* and *G*/*L* varied with whisker column identity, but *C*/*L* did not (R^2^=0.71,p<0.0001 for *m*/*L*; R^2^=0.56, p< p<0.0001 for *G*/*L*; R^2^=0.09, p=0.05 for *C*/*L*; Student’s two-sided t-test). However, *C* varied directly with column (Fig. S2). The systematic changes of *m*/*L* and *G*/*L* with column position suggested that follicle rotation might also vary systematically across the array. Therefore, we next aimed to simulate the effects of *G, L, m*, and *C* on follicle rotation.

### 2.2. A uniform motor command causes follicles in all columns to rotate through approximately equal angles

In general, muscle contraction depends on pennation and fiber type, but rat intrinsic whisker muscles are not pennate ^43^, and ∼90% of sling fibers are type IIb/d ^44^. Thus, sling muscle length alone determines contraction speed, and changes in length were used to simulate follicle rotation.

We began by defining the three angles of the *GLm* triangle − *θ*_*G*_, *θ*_*L*_, and *θ* _*m*_ − and the “motor angle” *θ* _*motor*_ (Fig. 2A). The motor angle represents the whisker’s angle relative to the skin for any *G, L*, and *m* combination in each follicle’s local reference frame. Next, we computed the average *GLm* triangle across animals for each whisker column, normalized by *L* (Fig. 2B). If these triangles were geometrically similar between columns, follicle rotation would be as well. However, plotting *θ*_*G*_, *θ*_*L*_, and *θ* _*m*_ as functions of column position (Fig. 2C) shows otherwise: the angle *θ*_*G*_ is relatively constant across columns, while *θ*_*L*_ decreases and *θ* _*m*_ increases.

**Figure 2.**
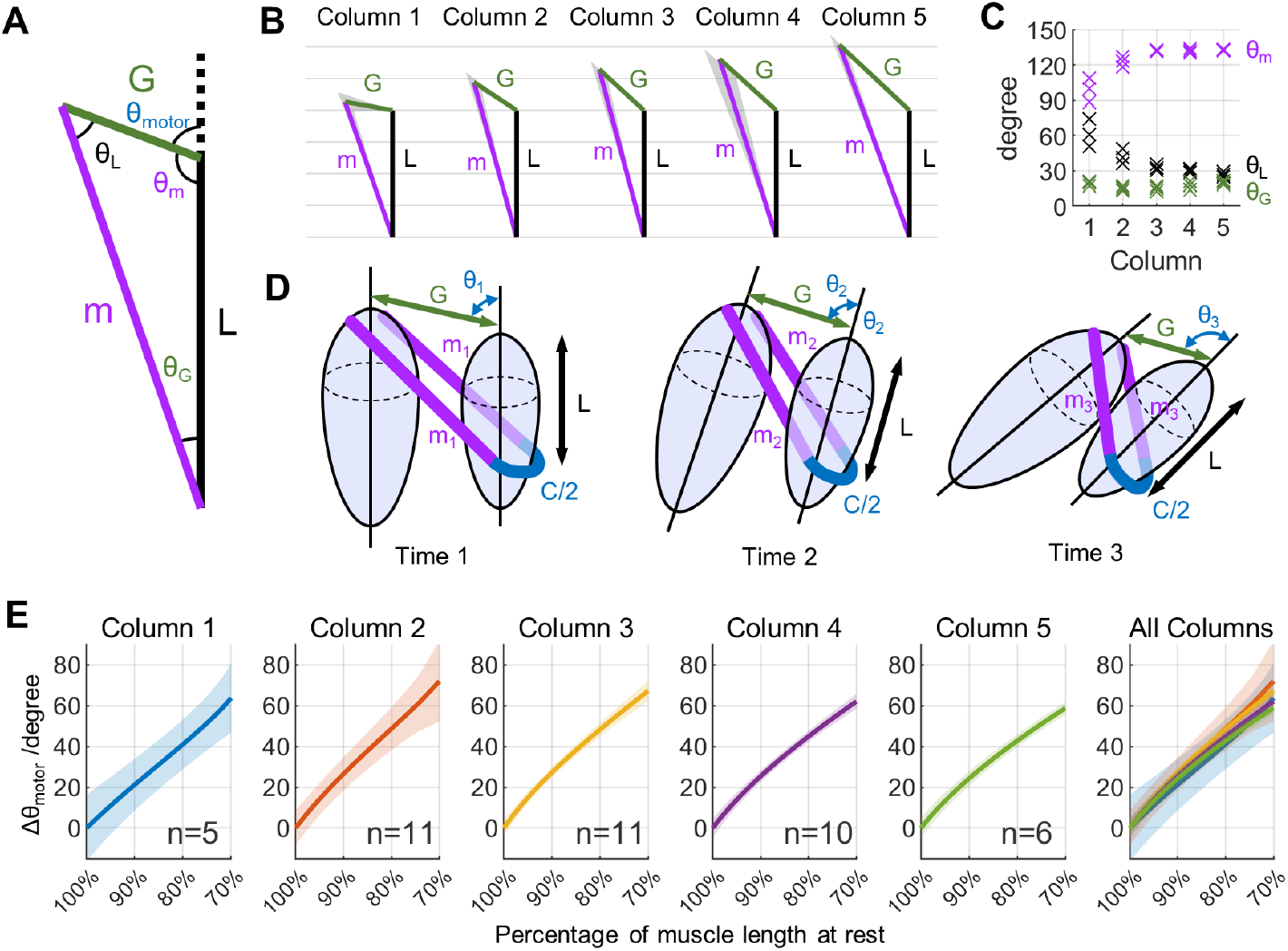
Simulations of follicle protraction from measured parameters. **(A)** Schematic of angles relevant to simulating whisker protraction. The angle *θ* _*motor*_ is the angle that the whisker forms relative to the skin, for any combination of *G, L*, and *m*. **(B)** Each subplot shows the *GLm* triangle averaged across animals and across all whiskers within that column. Values of *L, G*, and *m* are normalized by *L*. Shaded patches around the lines *G* and *m* indicate standard deviations. **(C)** Plotting the three angles within the *GLm* triangle as a function of column identity indicates that the triangles are not similar across columns. Each point represents data averaged from a single rat. **(D)** Simulated protractions based on the *GLm* triangles shown in (B) and follicle circumference. Total muscle length decreases by a fixed percentage each step, updating m and θ. **(E)** Simulations of whisker protraction show that equal changes in muscle length cause follicles in all columns to protract through approximately equal angles. The simulation stops when the muscle is at 70% of its resting length. The value n indicates the number of follcles averaged in each column (see Table S1)

We used the *GLm* triangles for each of the 43 reconstructed follicles to simulate the follicle rotations that would occur if all intrinsic muscles contracted an equal percentage. Simulation details are in *Methods*, but briefly, the simulation consisted of each rostro-caudal pair of follicles moving in a shared plane (Fig. 2D). The starting angle (*θ*_1_) of each simulated whisker was set to the initial value of *θ* _*motor*_ determined for that follicle, and initial muscle length set to 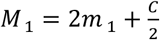. During each timestep of the simulation, *M* was reduced by a percentage and the value for muscle arm length *m* recomputed, while keeping *C* constant. Each follicle was then rotated about its apex to match the new value for *m*, and resulting whisker angles calculated from the law of cosines. An important characteristic of the simulation is that follicle rotation depends both on *m*, which shortens during contraction, and on *C*, which remains fixed.

Simulation results, shown in Fig. 2E, revealed that follicles in all columns rotate through approximately equal angles in their local reference frames for a given percent contraction. In other words, if the identical motor command is delivered to a rostral and caudal follicle, they will rotate through approximately equal angles. Changes in whisker protraction angles are not correlated with column at any muscle contraction level up to 70% (p > 0.05), and show only weak correlation at 70% (p = 0.0463; two-sided t-test), indicating similar slopes across columns within measurement error. The simulation shows unphysiological results once muscle length falls below ∼70% of resting, consistent with prior models of rodent facial muscles ^10^ and sarcomere contraction limits ^45^.

A sensitivity analysis (*Methods; Fig. S3*) established how much each parameter (*G, L, m, C*) affected the slopes in Fig. 2E. When comparing *GLm* triangles of different sizes, Δ*m*/*m* has the largest effect on the slope, followed by Δ*G*/*G* and Δ*L*/*L*. In contrast, follicle rotation is relatively insensitive to variations in circumference, suggesting that it may be shaped by selective pressure for sensory acquisition rather than motor control; e.g., maximizing circumfrence would maximize mechanoreceptor density.

### 2.3. Consequences of equal follicle rotation for 3D whisking profiles and the rat’s sensory space

So far, results have demonstrated that all follicles rotate through nearly-equal angles during a uniform motor command, in which sling muscles contract by an equal percent. These rotations (motor angles) are computed within the local, two dimensional (2D) planar coordinate system of the follicle’s specific *GLm* triangle. We simulated how these 2D (planar) rotations affect the rat’s 3D whisking trajectories as seen from above, and how they affect the rat’s 3D sensory space (Fig. 3; Video S2).

**Figure 3.**
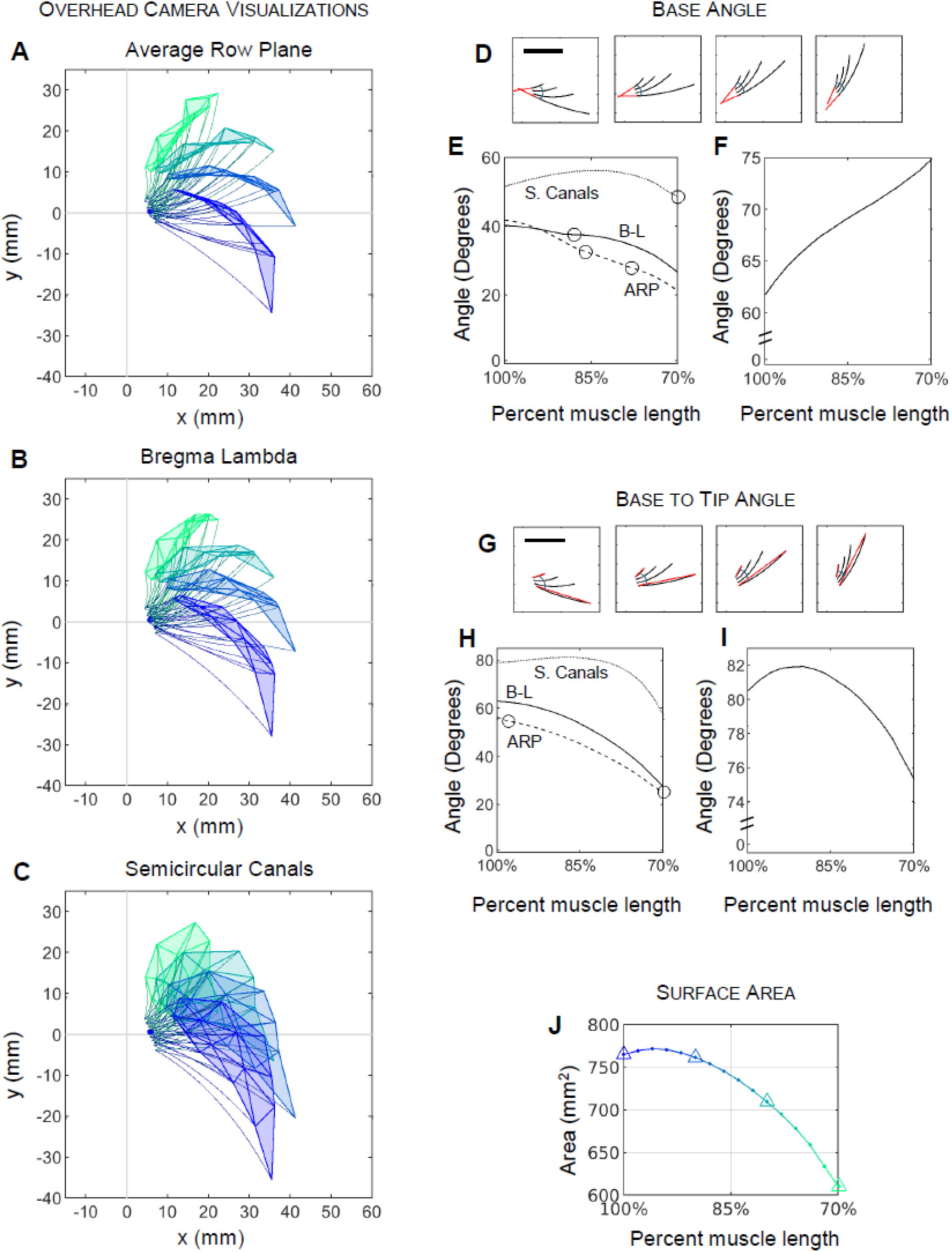
Simulated whisker kinematics during uniform contraction of the sling muscles. (ABC) Visualization of the rat’s sensory space during a uniform motor command. Three panels show whisker tip positions projected onto three horizontal planes, each corresponding to progressively greater downward pitch of the rat’s head: average row plane, bregma-lambda plane, and semicircular canal plane. Colors shift from purple, blue, cyan, to green as percent muscle length decreases from 100%, 90%, 80%, 70%. **(D)** Projection of the “base angle” for C-row whiskers onto the bregma-lambda plane at four protraction timesteps. **(E)** Projected base angle magnitude as a function of percent muscle length, for three horizontal planes. **(F)** Corresponding 3D base angle magnitude shows monotonic increase during protraction. **(G)** Projection of “base-to-tip angle” for C-row whiskers onto the bregma-lambda plane at four protraction timesteps. **(H)** Projected base-to-tip angle magnitude as a function of muscle length across three planes. **(I)** Corresponding 3D base-to-tip vector magnitude peaks at approximately 90% muscle length. **(J)** Surface area enclosed by whisker tips as a function of percent muscle length, peaking early and then decreasing continuously.

In these simulations, 15 of the 17 whiskers shown in Table S2 were initialized to their 3D resting positions and orientations established in previous work ^42^. The two whiskers in Column 1 were excluded due to the large error bars observed in Fig. 2E and the results of the sensitivity analysis. For each percent muscle contraction, values of *θ*_*motor*_ were calculated for the whiskers in columns 2 − 5, exactly as in Fig. 2E. These values of *θ*_*motor*_ were then projected into a chosen horizontal plane, and the resulting rotation within the plane was defined as the whisker’s “protraction angle.” After computing the protraction angle, both roll and elevation were incorporated according to previously-established kinematic equations ^46^.

Fig. 3A illustrates simulation results for three choices of horizontal plane, each corresponding to progressively greater downward pitch of the rat’s head: the average row plane, the bregma-lambda plane, and the semicircular canal plane. These visualizations represent what would be observed by an overhead camera oriented parallel to each plane during uniform sling muscle contraction. For all three planes, whiskers converge during protraction, resulting in a reduction in the surface area enclosed by the whisker tips near peak protraction (green polygon) relative to the area enclosed at rest (purple polygon). Comparing across the three panels, it is clear that increasing downward pitch of the rat’s head results in greater apparent overlap in whisker-tip locations in overhead views.

To explore these ideas further, we quantified the array “base angle,” defined as the angular difference in 3D orientation between the two most divergent whiskers as they emerge from the follicles. The orientation of each whisker was computed as a vector extending from its basepoint to a point located 1% along its length. For each percent muscle contraction, the 3D angle between every whisker pair was calculated, and the array base angle was defined as the maximum angular separation at each step. The array base angle was then projected into selected planes to compute its 2D form.

Fig. 3B depicts the array base angle for the C-row whiskers, projected onto the bregma-lambda plane, at four discrete timesteps during a protraction. In this 2D projection—corresponding to an overhead camera view—the whiskers appear to converge. This trend is quantified in Fig. 3C, where the projected base angle magnitude decreases from approximately 40 to 30 degrees with increasing muscle contraction. Similar results are observed for projections into the average row and semicircular canal planes (dashed and dotted lines). In contrast, the 3D analysis (Fig. 3D) reveals that the true base angle magnitude increases from 62 degrees to 75 degrees over the same interval. This discrepancy arises because the whiskers diverge in the dorsoventral direction during a protraction, expanding the rat’s sensory coverage.

A similar analysis can be performed for the array “base-to-tip” angle, defined as the angular difference between the 3D orientations of the two most divergent whiskers, measured from their basepoints to their tips (Fig. 3E). In the bregma-lambda projection, array base-to-tip angle magnitude decreases monotonically during protraction (Fig. 3F), with similar trends observed in the average row and semicircular canal planes. In contrast, the corresponding 3D analysis reveals a non-monotonic profile: the magnitude initially increases, peaks at approximately 90% muscle length, and then decreases (Fig. 3G).

Finally, plotting the surface area enclosed by the whisker tips as a function of percent muscle length (Fig. 3H) closely mirrors the trends observed for the array base-to-tip angle. Surface area reaches its maximum shortly after protraction onset and then decreases steadily as muscle contraction progresses. A previous study analyzing multiple bouts of non-contact whisking behavior reported an approximately linear relationship between whisker-tip surface area and whisker spread ^47^. The present simulations show a similar near-linear relationship once muscle length falls below 90% of resting. This result suggests that during fine tactile exploration, rats may avoid fully retracting their whiskers, maintaining them instead within a range where uniform motor commands produce predictable, linear changes in the sensory space.

Summarizing, Fig. 3 demonstrates that when all sling muscles contract by the same percentage, the angular spread of whiskers at their base increases monotonically throughout protraction. In contrast, the maximum distance between whisker tips occurs at approximately 90% of the resting muscle length, after which the spatial separation—and thus spatial resolution as measured by tip distance—decreases monotonically. A uniform motor command serves to linearly position all whiskers in rostral regions near the snout, potentially reducing cognitive demands. Future studies may explore roles of additional extrinsic facial muscles, which were not included in the present simulations ^4-7,9,10,12,38,48^.

## Resource availability

## Lead contact

Requests for further information and resources should be directed to, and will be fulfilled by, the lead contact, Mitra Hartmann

## Materials availability

This study did not generate new unique reagents.

## Data and code availability

- All experimental data reported in this paper will be deposited in a publicly repository on the Open Science Framework
- Any additional information required to reanalyze the data reported in this paper is available from the lead contact upon request.

## 3. STAR*Methods

### 3.1. Key resources table

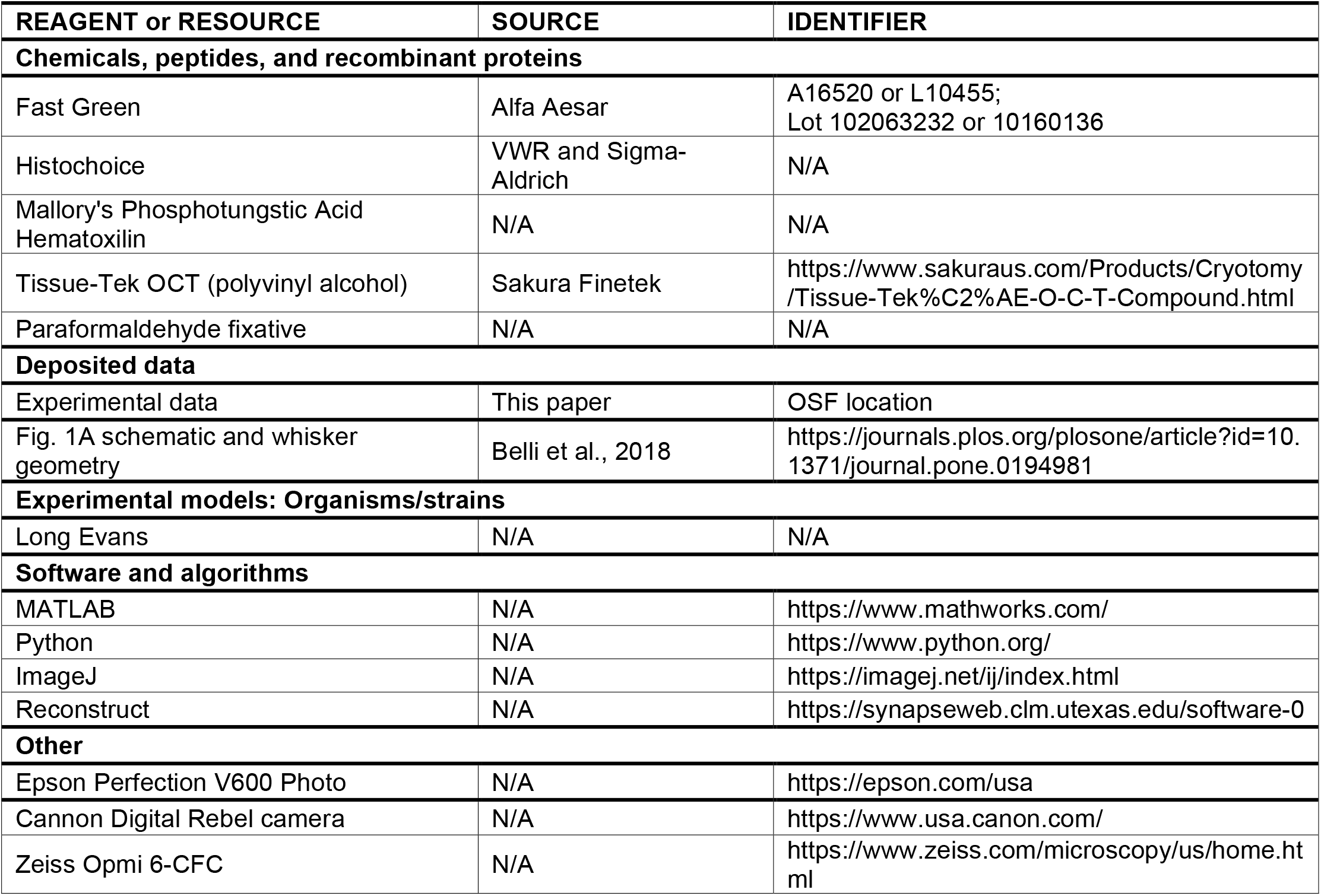

### 3.2 Experimental model and study participant details

All procedures were approved in advance by the Animal Care and Use Committee of Northwestern University.

The present study was based on anatomical reconstruction of the mystacial pads of three Long Evans rats. Two rats were females (6 − 18 months), and the third was male (3 months).

### 3.3. Method details

#### 3.3.1. Perfusion, dissection, embedding, sectioning, and staining

Our approach was carefully designed to address three separate problems often associated with histological analyses of gross anatomy. First, mechanical distortion of the soft tissue surrounding the follicles was minimized by sectioning entire pads without flattening, and without dissecting individual rows or follicles. Second, we minimized uneven muscle contraction during fixation by using a non-formaldehyde-based fixative, and checked for any remaining unevenness by measuring relative sarcomere lengths to assess non-uniform stretch or contraction across the pad. Third, we minimized tissue shrinkage and distortion during specimen preparation by using frozen sections that required less aggressive dehydration than paraffin or plastic sections, and by very quickly freezing the tissue on an aluminum block cooled with liquid nitrogen. Each of these procedures is described in more detail below.

After use in unrelated electrophysiology experiments, rats were perfused with 1x phosphate buffered saline solution (PBS) with 10 units/mL heparin and then with HistoChoice™. HistoChoice™ was used in place of formaldehyde because the sling muscles are type II fibers, and thus susceptible to formaldehyde-induced contraction. To dissect the mystacial pad, scissors were used to free the soft tissue of the snout from the underlying bone in two large flaps, each containing all macrovibrissae with a few mm boundary on all sides. The two flaps were detached from the underlying bone by gently peeling up the tissue flap and using microscissors to cut the anchoring connective tissue. After dissection, the tissue was placed in 100% HistoChoice™ overnight. After 24 hours, tissue was sequentially cryoprotected in 10%, 20%, and 30% sucrose in PBS until the tissue rested on the bottom of the vial, indicating that osmotic pressure had equalized.

After the tissue had sunk in the 30% sucrose solution, it was flash-frozen in Optimal Cutting Temperature compound (Tissue-Tek OCT, Sakura Finetek) on a level aluminum block partially submerged in liquid nitrogen. We ensured that the block was temperature equilibrated by visually confirming that boiling had stopped at the liquid/block interface. This procedure ensured that one face of the tissue was in contact with a surface very near -195° C, allowing a fast-moving unidirectional front of ice crystal formation to pass through the tissue. This quick unidirectional freezing is believed to generate very small crystals and minimize compression or stretching that would warp the tissue. Tissue was then sectioned at either 20 microns (pads 1 and 2) or 50 microns (pad 3) on an upright freezing microtome and mounted on gelatin-coated slides.

Tissue sections were fixed to gelatin coated slides using a 4% paraformaldehyde solution (PFA) for 15 min, permeabilized with acetone for 5 min, washed, bleached, stained in Mallory’s Phosphotungstic Acid Hematoxilin (PTAH), washed and dehydrated, stained in 0.1% Fast Green (FG) in ethanol, washed, cleared, and coverslipped. FG stains collagen blue-green, and PTAH stains muscle striations purple-blue and many tissues (including collagen) various shades of red-pink. When we double-stained for collagen and muscle the pink PTAH pigments were washed out with ethanol and the collagen was re-stained with FG to achieve darker and more distinct color.

To obtain the five images shown in Fig. 1C, photomicrographs of whole pad slices were taken with a Cannon Digital Rebel camera, mounted on a Zeiss Opmi 6-CFC dissecting scope, photographed at 8x magnification. These images were manually stacked and aligned solely for the purposes of illustration; red asterisks were added to visualize the same follicle between sections.

#### 3.3.2. Sarcomere length measurement to control mechanical skewing of follicle long axis

We performed a set of experiments to control for mechanical skewing of the long axis of the follicle. Specifically, to confirm that all sling muscles across the pad had contracted approximately the same percentage during fixation, we compared sarcomere length in experimental and deliberately-stretched positive control pads.

Sarcomere length was analyzed in six pads. Four of these pads came from animals that had been perfused and fixed while their whiskers remained in a relaxed position. Two of these four pads corresponded to Experimental Pads 1 and 2 from the *Results* section. Sarcomere length was not analyzed for Pad 3. The remaining two pads underwent the identical procedures as Experimental Pads 1 and 2 and are referred to as “Fixed Control” (FIX) pads. During fixation, the sling muscles in all four of these pads—the experimental pads and the two FIX pads— will tend to contract slightly, causing the whiskers to protract and the sarcomere width to decrease from its resting value. If the sling-muscle contraction is relatively uniform across the pad, we expect the sarcomere width to be nearly-constant across the array for these four pads.

Two additional pads served as positive controls and were obtained from animals that had been perfused and fixed with a subset of their whiskers deliberately taped in a retracted position. Taping whiskers into retracted positions will cause the sling muscles to stretch and the sarcomere width to increase from its resting value. Because only some of the whiskers were taped, we expect slightly larger average sarcomere widths and much higher variability in sarcomere widths across the array. These positive controls were designated as “stretched” (STR) pads.

Consistent with expectations (Fig. S1), samples of sarcomere widths taken across the experimental pads and the FIX showed relatively low average widths and low variability, indicating that sling muscle contraction was relatively uniform across the pad. In contrast, the STR pads exhibited larger average widths (2.7*μm*) and higher variability.

**Figure S1.**
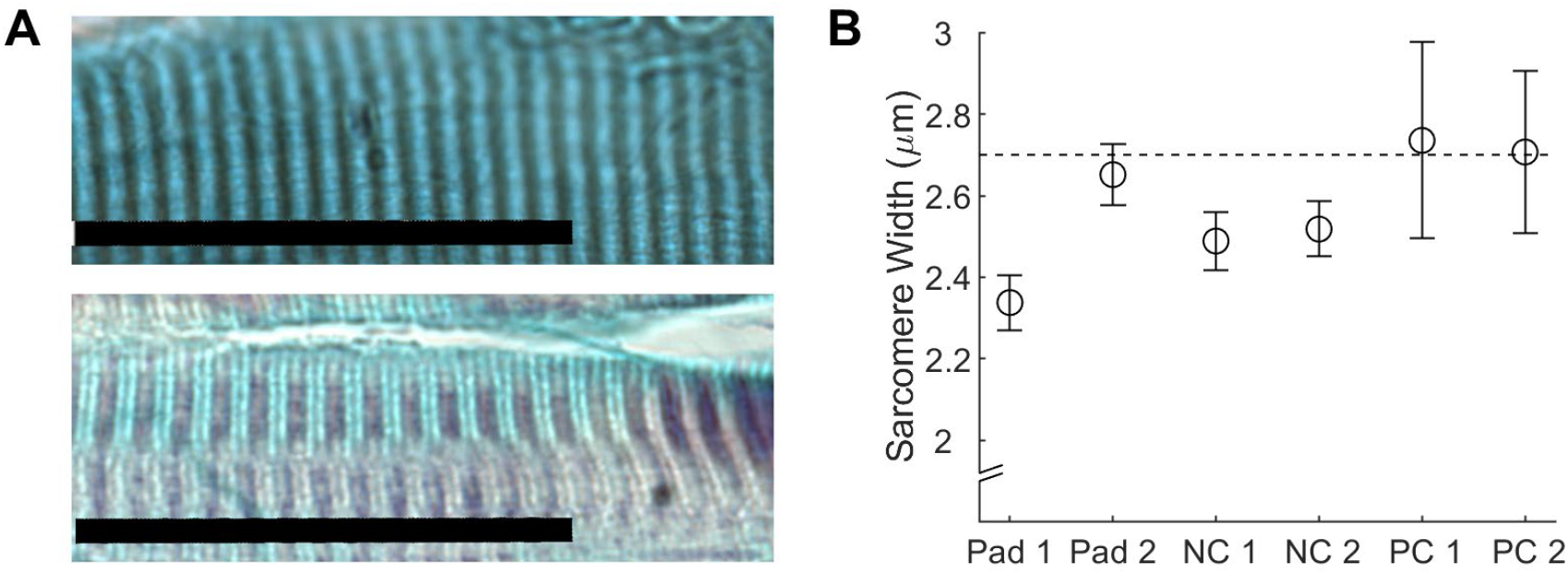
Analysis of sarcomere stretch controlled for whisker perturbation, non-uniform muscle contractions across the pad, and mechanical deformations of the pad. Experimental pads and the two negative control, fixed (FIX) pads exhibit lower mean sarcomere length and lower variance than the two positive control, deliberately stretched (STR) pads. **(A)** Example images of sarcomeres from a sling muscle in one of the FIX pads (upper panel), and one of the STR pads (lower panel). Sarcomeres are visibly narrower in the FIX pad than in the STR pad. Scale bars: 40 microns. Both images were taken at 40x. **(B)** Distribution of sarcomere lengths in two experimental pads, two FIX pads and two STR pads. Within each experimental and FIX pad, sarcomere lengths show relatively low variability (average SD is 0.07 *μm*) and are all close to 2.5 *μm*. In contrast, the two STR pads have a larger average sarcomere length (2.7*μm*; dotted line) and greater variability (± 0.24 *μm*). Related to Fig. 1 and the STAR*Methods.

#### 3.2.3. Scanning, image processing and registration, and reconstruction of follicle shape

The sectioning and staining process generated a set of slides for each mystacial pad, with each slide containing multiple mystacial pad sections. After drying, each slide was scanned at high resolution (6,400 dpi) on a flatbed scanner (Epson Perfection V600 Photo). The slide was positioned so that only a single section was placed in the scanner’s region of interest (ROI). Sometimes the ROI included the edge of a section adjacent to the one centered in the ROI.

After scanning, several image processing steps were performed within the software *ImageJ* ^49^. First, images were converted to 8-bit grayscale, and the edges of any adjacent sections were manually cropped out, so that each image contained only one complete section. Second, we visually identified and excluded poor quality images. Exclusion criteria included sections that did not contain follicles; sections at the most dorsal regions of the follicle that did not contain the follicle wall; sections that were folded so that follicles were obscured; and sections that were stretched so that the spacing between the follicles was obviously skewed. Third, using a reference slide chosen to be near the middle of the stack (so that all follicles were included), we ran the *ImageJ* plugin “Register Virtual Stack Slices” (RVS) to perform an initial alignment of all follicles. Alignment was then visually evaluated by scrolling through the stack using the mouse wheel, and single images with poor alignment were manually removed.

After initial alignment, standard Python image processing tools were used to automatically trace the outline of each follicle in each section and fit an ellipse to each follicle border. Manual checks were needed to remove false positives, track missed follicles, retrack inaccurate follicles, and to label follicles in select sections. These arrays of ellipses were then combined into z-stacks, with each follicle forming an irregular, slim volume (approximately cylindrical), usually slightly truncated at the apex and/or base depending on imaging and slicing quality.

Consistent with previous studies ^39,40^, initial reconstructions demonstrated that the follicles often curved or deformed near the base. We have a two-part explanation for this effect. First, the collagen wall of the follicle is thick and stiff in the upper (superficial) two-thirds of the follicle, but becomes much thinner in the deeper third, near the follicle base ^7^. In addition, the keratin that forms the whisker shaft becomes less dense near the follicle base, and the presence of live keratinocytes in this region further increases flexibility ^50^. Thus overall, the lower third of the follicle is much more flexible than the upper two-thirds. Second, because animals were anesthetized when they were euthanized, their blood pressure was low. Thus the blood sinus within the follicle was not enlarged, and fluid pressure would not have helped to stiffen the follicle near its base as may occur in the awake animal ^35^. Combined, these two effects could explain the observed curve near the base of each follicle.

To adjust for this deformation and ensure that the reconstructed follicles accurately represented their position and orientation within the mystacial pad required three steps.

First, follicle centers across the entire pad were aligned in accordance with the general bounds of whisker emergence angles previously established in anatomical reconstructions ^42^. To do this, we chose a “reference” image from the middle of the z-stack, being careful to ensure that all follicles were clearly visible. At the center of each follicle section, we placed an imaginary line oriented to match the emergence angles of the corresponding whisker. The group of imaginary lines together formed a “template follicle array”, which gave us a set of reference points at each z-value for the predicted location of each follicle in each section (image). The follicles in each section were then aligned as a whole to the template array, minimizing the summed distance between the center of each follicle cross-section and its position predicted by the template follicle array at that depth.

Second, each follicle was manually cleaned to remove tracking that did not accurately capture the follicle cross-section. For example, if only a fraction of a follicle circumference was tracked, its cross-section would often have a very odd shape Most of these outliers were superficial, representing regions after the whisker had exited from inside the follicle (i.e., between the follicle apex and the skin surface). After these outliers had been removed, each tracked follicle cross-section was fit with an ellipse.

Third, each follicle was aligned on its own principal axis of a stack of ellipse centroids. Although the alignment in the first step ensured that all follicles were well matched to the emergence angles, the cross-sections within individual follicles were not aligned smoothly. To reconstruct individual follicles, a 3D follicle body was generated from the stack of clean ellipses, up-sampled at every micron to increase the resolution between cross-sections, and then smoothed using a sliding window of 10x the original slice resolution. Each follicle was then resliced perpendicular to its principal axis based on smoothed, high-res follicle body. The follicle outline in each resliced section was used to estimate the follicle circumference at that depth.

For all three pads, we found that the slice angle was too severe to accurately reconstruct follicles from the A-row, the B1 and E1 follicles, and follicles rostral to column 5. These follicles were therefore eliminated. In addition, we eliminated select follicles with poor reconstruction quality from individual pads. Table S1 lists all reconstructed follicles from all three pads: 16 follicles were obtained from Pad 1, 11 follicles from Pad 2, and 16 follicles from Pad 3, for a total of 43 follicles. Relationships amongst all reconstructed parameters are shown in Fig. S2. Statistical analysis of these parameters demonstrated that *L* and *m* decreases linearly with column identity, whereas *G* shows no correlation with column identity.

**Table S1.** A list of all follicle/muscles reconstructed in the present study. We were able to reconstruct most follicles/muscles in the central mystacial pad (“pad”) region. Follicles in the dorsal and rostral regions of the mystacial pad were sliced at severe angles and thus could not be reconstructed accurately. **“**N/A” indicates that the whisker is not associated with a standard sling muscle, and was not analyzed in the present study. “None” (red) indicates the whisker is associated with a sling muscle, but no reconstructions were obtained. The region of reconstruction is in bold text. A horizontal dash (“—”) indicates that a reconstruction could not be obtained for the indicated pad(s). All pads are from Long-Evans rats. Pads 1 and Pad 2 are from 6 − 18 mo. female rats; Pad 3 is from a ∼3 mo. male rat. Related to Fig. 1 and the STAR*Methods.

|  | “Greek” | Col. 1 | Col. 2 | Col. 3 | Col. 4 | Col. 5 | Col. 6 | Col. 7 | Total |
| --- | --- | --- | --- | --- | --- | --- | --- | --- | --- |
| **A row** | N/A | *none* | *none* | *none* | *none* | N/A | N/A | N/A | 0 |
| **B row** | N/A | *none* | **Pad 1**<br>—<br>**Pad 3** | **Pad 1**<br>—<br>**Pad 3** | **Pad 1**<br>—<br>— | *none* | N/A | N/A | 5 |
| **C row** | N/A | **Pad 1**<br>—<br>**Pad 3** | **Pad 1**<br>**Pad 2**<br>**Pad 3** | **Pad 1**<br>**Pad 2**<br>**Pad 3** | **Pad 1**<br>**Pad 2**<br>**Pad 3** | **Pad 1**<br>—<br>**Pad 3** | *none* | *none* | 13 |
| **D row** | N/A | **Pad 1**<br>**Pad 2**<br>**Pad 3** | **Pad 1**<br>**Pad 2**<br>**Pad 3** | **Pad 1**<br>**Pad 2**<br>**Pad 3** | **Pad 1**<br>**Pad 2**<br>**Pad 3** | **Pad 1**<br>**Pad 2**<br>**Pad 3** | *none* | *none* | 17 |
| **E row** | N/A | *none* | **Pad 1**<br>**Pad 2**<br>**Pad 3** | **Pad 1**<br>**Pad 2**<br>**Pad 3** | **Pad 1**<br>**Pad 2**<br>**Pad 3** | —<br>—<br>**Pad 3** | *none* | *none* | 10 |
| **Total** | N/A | 5 | 11 | 11 | 10 | 6 | 0 | 0 | **43** |

**Figure S2.**
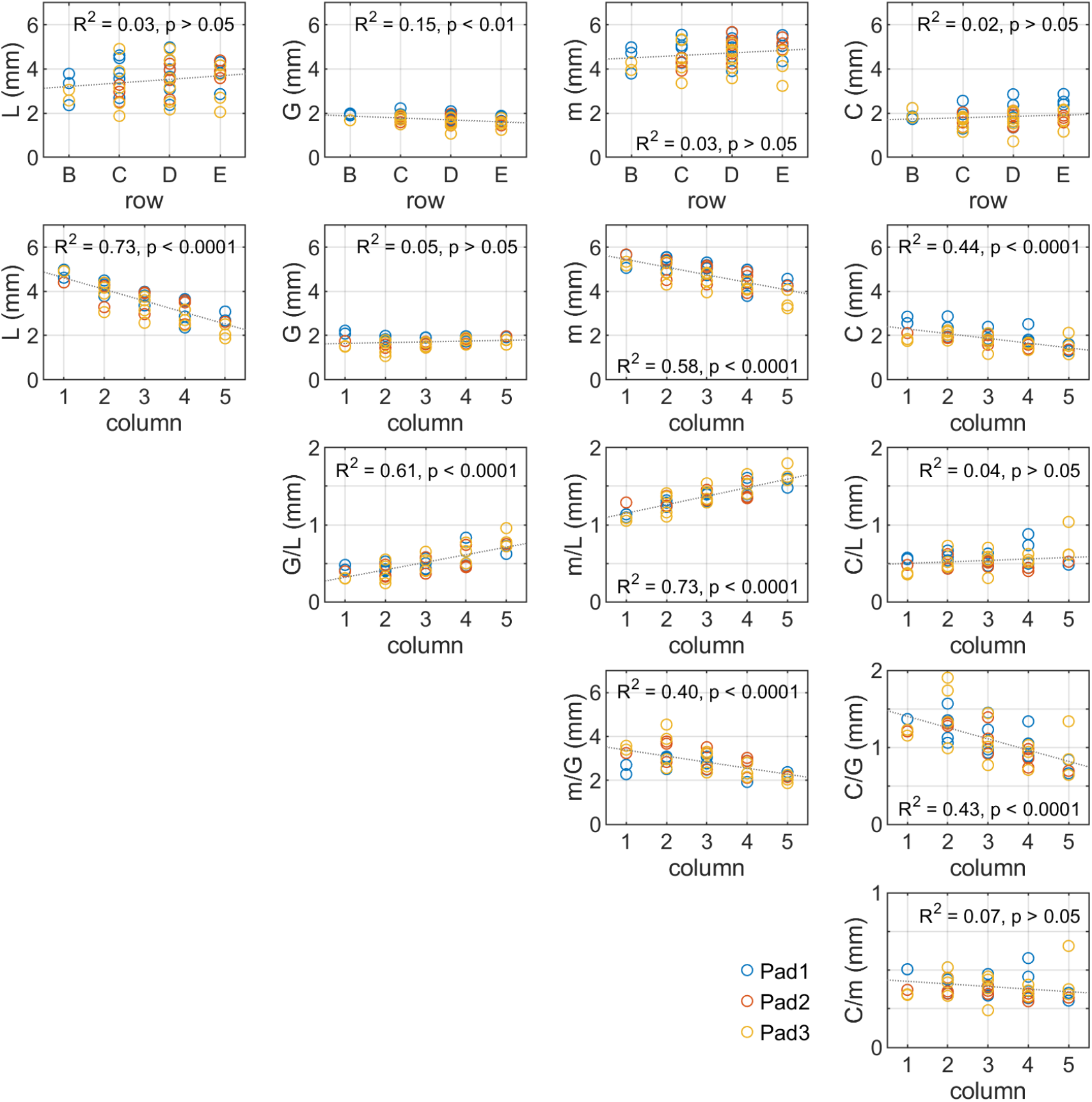
Relationships between reconstructed parameters. Blue: pad 1; pink: pad 2; yellow: pad 3. Related to Fig. 1 and STAR*Methods. The significance of each fit (two-sided Student’s t-test) is indicated in each panel.

### 3.4. Finding muscle attachment points

We searched through the anatomical slide sections from lateral to medial, and, for each follicle pair within a row, identified the first section in which the sling muscle fully wrapped around the rostral follicle. The section number was noted and the corresponding location was found in each resliced follicle. With that contact point determined, the circumference C of the follicle at the location of the muscle attachment point could then be determined in the (resliced) sections. The lever length L was computed as the distance between the (x, y, z) follicle apex and the muscle-follicle contact point. The gap G was computed as the distance between row-wise adjacent (x, y, z) follicle apices. The muscle arm length m was computed as the average length of the two common tangents between the two tracked follicle outlines, at the levels of the follicle apex and the muscle attachment point.

### 3.4. Quantification and statistical analysis

#### 3.4.1 Computing the geometry of follicle rotation and whisker protraction

The follicle rotation simulation consisted of a rostro-caudal pair of follicles, with the rostral follicle moving in their shared plane, as illustrated in Fig. 1D. For each pair of follicles the simulation was initialized based on measured values for the *GLm* triangle and the follicle circumference. The initial motor angle *θ*_1_ is calculated from measurements of L, G, and m^1^. The full sling muscle length when at rest is given by 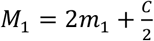, where *C* is the follicle circumference.

At each time step *t* in the simulated simulation, the resting length of the intrinsic muscle *M* was contracted by a given percentage *x* (*M*_*t*_ = *M*_*t*−1_ − *XX*% of *M*_1_), and the value of the muscle arm *m*_*t*_ was recomputed as:

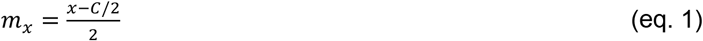

The new follicle motor angle *θ*_*x*_ was calculated using the law of cosines:

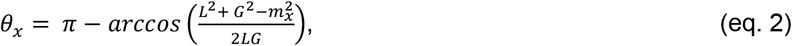

where *x* is a fraction between 0 and 1, representing the percentage of the resting muscle arm length that the muscle was contracted to.

It is important to note that follicle rotation angles, computed in the coordinate system local to the GLm triangle, are not the same as whisker protraction angles, computed within a single head-centered coordinate system. For each follicle, the value of *θ* _*motor*_ is defined within the plane of its GLm triangle: this angle represents rotation of the follicle relative to its local skin surface. If the GLm triangle of particular follicle happens to lie in the horizontal plane (defined with respect to the head), then the follicle motor angle will be identical to the whisker protraction angle, but otherwise these two angles will be different. If this claim seems unintuitive, it may be helpful to visualize the lines that connect each row-wise pair of follicles: each of these lines has a slightly different 3D orientation.

#### 3.4.2. Sensitivity analysis

Our goal is to determine how variability in each parameter *L, G, m, C*, will affect the slopes 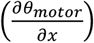 of the curves shown in Fig. 2E. We start from the two equations previously used to compute the geometry of follicle rotation. To simplify notation, we drop the subscript “motor,” and define *θ* ≡ *θ*_*motor*_. Because we are interested in the variability in *L, G, m*, and *C*, the motor angle *θ* is expressed as a function of these quantities by substituting equation 1 into equation 2:

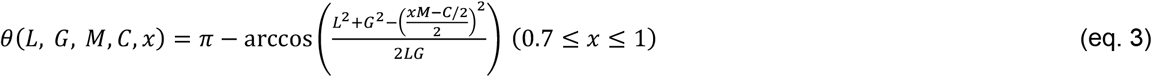

where the constraint (0.7 ≤ *x* ≤ 1) limits use of equation 3 to a biologically-plausible range.

Substituting 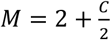 yields

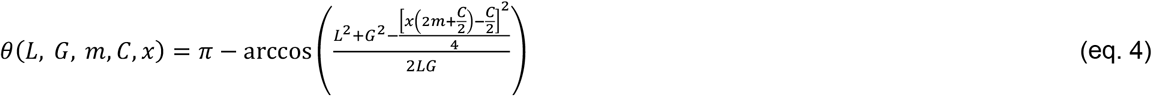

Simplifying:

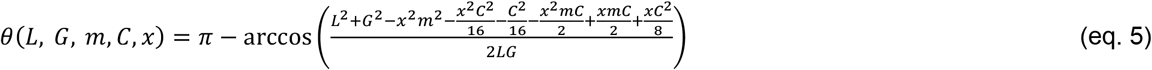

To simplify equation 5, we noted that *θ* for the experimental dataset tended be close to a right angle (Fig. S3A) In other words, the argument of the arccosine tended to vary around zero, with the notable exception of the column 1 follicles. As shown in Fig. S3B, when all follicles (columns 1 − 5) were included, the argument ranged between -0.6890 and 0.8296. When column 1 was omitted, however, the upper value decreased to 0.5729. Fig. S3C further shows that the percent difference between the function arccosine and its Taylor series expansion 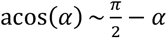 differs by less than 4% over the range for follicles in columns 2 − 5, thus supporting the use of a first-order Taylor series expansion for columns 2 − 5 for the remainder of the analysis

**Figure S3.**
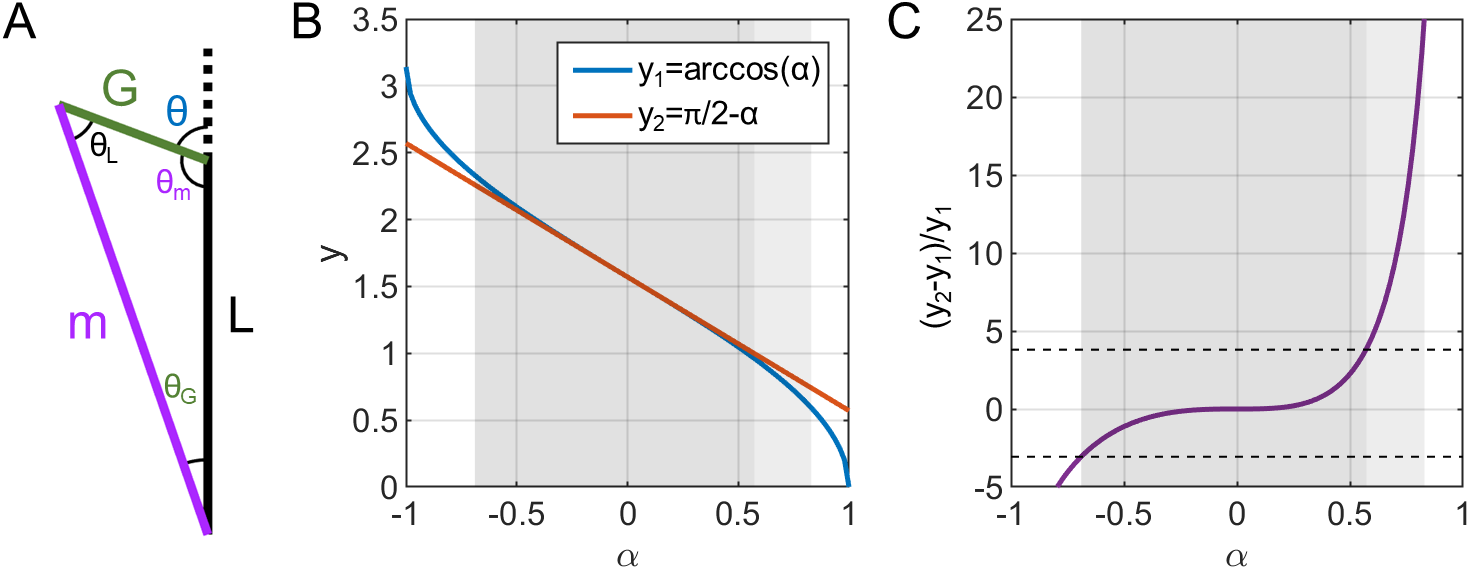
(A) The value of *θ* for most GLM triangles tended to be close to 90°, suggesting that a Taylor series expansion could be used to simplify equation 5. **(B)** The arccosine function (*y*_1_, blue) and its first-term Taylor series approximation (*y*_2_, orange). Experimental data showed that the GLM triangles across all columns had values of *θ* in the range between 0.6890 and 0.8296 (light gray shaded region). When column 1 whiskers were omitted, the range of *θ* values in the experimental data fell within the range 0.6890 and 0.5729 (dark gray rectangle). **(C)** Percentage difference between the arccosine function and its first-term Taylor series approximation, computed as 100 * (*y*_1_ − *y*_2_)/*y*_1_. When column 1 follicles are included (light gray shaded rectangle), the percentage difference is as large as ∼25%, an unacceptable approximation (light gray rectangle). Thus the sensitivity analysis performed here does not hold for the column 1 whiskers. When column 1 whiskers are excluded (dark gray rectangle), the approximation deviates from the true value only from -3.8% to 3.1%, values indicated by the dashed black horizontal lines.

Thus, using the Taylor series approximation 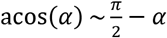 we can approximate equation 5 as:

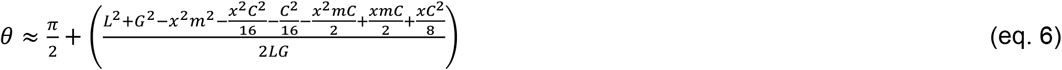

Recalling that our goal is to determine how 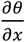 is affected by changes in *L, G, m*, and *C*, we denote 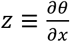 that:

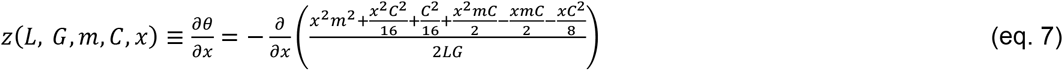

Simplifying:

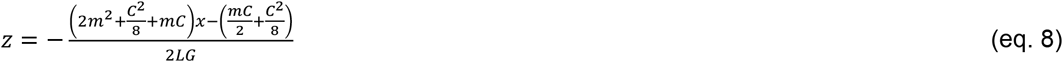

Noting that the numerator contains only *C, m* and *x*, while the denominator contains only *L* and *G*, we define

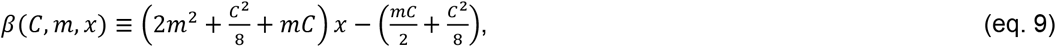

and now consider how each variable in turn affects 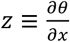

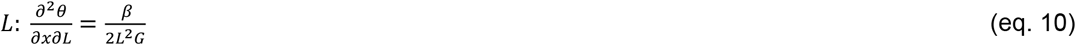

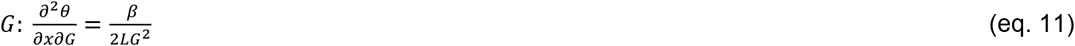

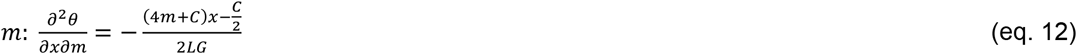

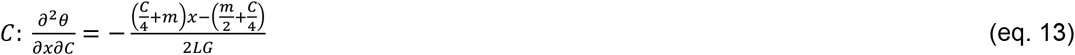

We now consider how each of *L, G, m*, and *C* affect *z* by considering the total differential for z:

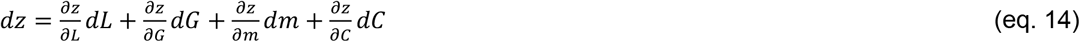

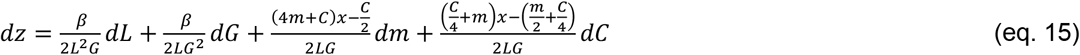

To examine how *dz* (in units of mm/radian) is affected by variability in each parameter, we substituted median values of *G* = 1.72 *mm, L* = 3.59 *mm, m* = 4.84 *mm, C* = 1.81 *mm, x* = 0.85 into equation 15 to arrive at (units omitted):

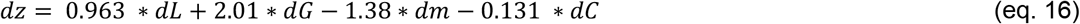

Or

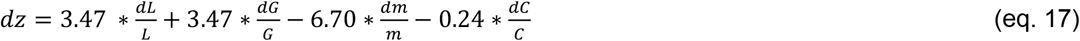

Equation 16 describes how variability in each parameter contributes to the variability in the slope. This equation reveals that future experiments should focus on minimizing measurement error in *G* as much as possible, as it has the largest influence on the slope.

Equation 17 describes how the slope is sensitive to a proportional change in these parameters. For GLm triangles of different sizes, *dm*/*m* contributes most to the change in the slope, followed by *dL*/*L* and *dG*/*G*. Because *m* is opposite in sign to *G* and *L* signs, proportional changes in *G, L, m*, and *C* are counterbalanced.

## Acknowledgments

This multi-year project was sequentially supported by NSF awards IOB-0846088, IOS-0818414, EFRI-0938007 and National Institutes of Health R01-NS116277 to MJZH. CS Bresee was supported in part by NSF IGERT: Integrative Research in Motor Control and Movement Grant DGE-0903637 and by NIH Grant T32 HD-057845. We thank Mr. Michael Penn for performing some of the histology, and Northwestern undergraduates Ariela Deleon, Aurora Greane, Delan Hao, Godson Osele, Liam Perreault, Pia Sanpitak, Jeannette Wu, and William Zeng for performing much of the follicle scanning and tracking. We thank Prof. Hayley Belli and Dr. Lucie Huet for writing components of the code and for many useful discussions.

## Supplementary Videos

Supplementary Video 1 rotates one of the three reconstructed follicle pads to visualize the follicles from multiple views.

Supplementary Video 2 shows the whisk of Fig. 3 from multiple viewpoints.

## Declaration of generative AI and AI-assisted technologies in the writing process

During the preparation of this work the authors used ChatGPT in order to explore alternative word choices and phrasing to try to improve scientific clarity. After using this tool/service, the authors reviewed and edited the content as needed and take full responsibility for the content of the publication.

